# Preclinical characterization of AT-03, a novel Serum Amyloid P fusion protein that demonstrates pan-amyloid binding and removal

**DOI:** 10.64898/2025.12.23.696051

**Authors:** Christophe Sirac, G. Roussine Codo, Arnaud Jaccard, Sébastien Bender, Gemma Martinez Rivas, Frank Bridoux, Marie Clavel, Camille Villesuzanne, Oussama Taoui, Claire Carrion, Gregory Bell, Spencer Guthrie, Suganya Selvarajah, Andrew Vick, Stephen J. Kennel, Tina Richey, Alan Stuckey, Manasi Balachandran, Angela D. Williams, Emily B. Martin, James S. Foster, Jonathan S. Wall

## Abstract

The systemic amyloidoses are progressive disorders caused by extracellular deposition of insoluble amyloid fibrils leading to organ dysfunction that often proves fatal. New therapeutics aiming at removing deposited amyloid are urgently needed to improve patient outcomes.

**Methods:** We developped AT-03 (originally called SAP-scFc), a fusion protein consisting of serum amyloid P-component, which binds all types of amyloid, linked to a single chain human IgG1 Fc domain. AT-03 binding to diverse types of amyloid and phagocytic activity were assessed both *in vitro* and *in vivo*. Therapeutic efficacy was evaluated in an AA mouse model.

**Results:** AT-03 bound with high potency to AL and ATTR human amyloid extracts. In murine models, intravenously administered AT-03 bound to AA, AL and AApoA2 amyloid, including in the heart. *Ex vivo* AT-03 opsonization induced phagocytosis of human AL extract by activated human THP-1 macrophages and enhanced *in vivo* phagocytosis in mice. A single intravenous injection of SAP-scFc induced a significant reduction of splenic amyloid in a murine model of AA amyloidosis.

**Conclusions:** AT-03 binds many amyloid types and can promote macrophage-mediated phagocytosis of the deposits. Thus, AT-03 is a promising novel therapeutic agent for the removal of systemic amyloid.

## Introduction

Amyloidosis is a severe, progressive, and often fatal protein misfolding and deposition disorder (1–3). There are approximately 18 different types of systemic amyloidosis, each caused by misfolding of a specific precursor protein, which then deposits in the extracellular space as insoluble amyloid fibrils (2, 4, 5). Accumulation of amyloid in organs leads to disruption of the surrounding tissue architecture and progressive organ dysfunction (6, 7). The most common types of systemic disease are transthyretin-associated (ATTR) amyloidosis, with an incidence of 16.6 per 100,000 in the US population of 65+ y of age and light chain-associated (AL) amyloidosis, with an incidence of 1.3 per 100,000 adults in the US (8). Other types of systemic amyloidosis are classified by the amyloidogenic precursor protein (1, 7). In ATTR, deposition of amyloid fibrils occurs most notably in nerve and cardiac tissue, resulting in varying severities of polyneuropathy and/or cardiomyopathy phenotypes (3, 9). The heterogeneity in clinical presentation and slowly progressive nature of these diseases frequently results in a delayed diagnosis or misdiagnosis (10).

The main determinants of morbidity and mortality in systemic amyloidosis are cardiac and renal involvement (11, 12). In cardiac amyloidosis, the accumulation of amyloid gives rise to cardiac arrythmias, restrictive cardiomyopathy, cardiac failure, and sudden cardiac death (3, 6, 9, 13)}. In patients with ATTR cardiomyopathy, the median survival from diagnosis is about 2 years for those with hereditary ATTR (ATTRv) and <4 years for those with wild-type ATTR (ATTRwt) (14). In patients with AL cardiac involvement, the overall median survival is around 2.6 years (15); after the onset of cardiac failure, the median survival drops to 6 months if left untreated (16). Renal involvement is also associated with significant morbidity, with a median survival after initiation of dialysis for amyloidosis-associated end stage renal disease of 8.2 months (17).

Current therapeutic approaches aim to reduce new amyloid deposition and slow disease progression by targeting the production of the amyloidogenic precursor protein (18–20). Approved therapies for use in patients with hereditary ATTR amyloidosis include transthyretin silencers (i.e., patisiran and inotersen) and tetramer stabilizers (i.e., tafamidis) (21). The only approved therapeutic regimen for AL amyloidosis is anti-plasma cell immunotherapy using daratumumab, a human CD38-targeting immunoglobulin G (IgG) monoclonal antibody (mAb) (22, 23). These therapies are most effective in the early stages of disease, but do not target the removal of existing amyloid fibril deposits (24). Clinical deterioration even after initiation of treatment is common, as many patients already have a significant accumulation of amyloid in vital organs at the time of diagnosis. Furthermore, while these therapies have made a significant impact in the treatment of ATTR and AL amyloidosis, there are no approved treatment options for patients with other types of systemic amyloidosis.

There is an urgent unmet need for novel therapies that effectively and safely promote amyloid removal of any type (pan-amyloid therapeutics), allowing restoration of organ function. A potential pan-amyloid targeting agent is serum amyloid P component (SAP), a pentameric protein that binds with high avidity to all types of amyloid (25–27). Dezamizumab, a fully humanized anti-SAP mAb, showed initial promise in two phase 1 trials in patients with systemic amyloidosis with liver, spleen, renal, or cardiac involvement (12, 28). In a subset of patients where major clearance of hepatic amyloid was demonstrated using SAP scintigraphy, a corresponding improvement in liver stiffness was also observed (12), providing preliminary evidence supporting amyloid-targeting immunotherapy to remove tissue amyloid. However, a phase 2 clinical trial showed no significant reduction in cardiac amyloid burden possibly due to the low uptake of dezamizumab in cardiac amyloid. The trial was subsequently terminated due to a serious adverse event (29).

AT-03 is a novel fusion protein consisting of SAP linked to a single chain human IgG1 fragment crystallizable (Fc) domain. Unlike dezamizumab, which requires prior and sustained depletion of circulating SAP in order to target amyloid-bound SAP (28), AT-03 exploits the pan-amyloid reactivity of SAP to deliver Fc domains to amyloid, which can then stimulate macrophage-mediated clearance of the deposits. The aim of this study was to characterize the preclinical profile of AT-03, including binding to human amyloid extracts, the ability to promote macrophage-mediated phagocytosis and clearing of amyloid, as well as pharmacokinetic behavior in non-human primates.

## Materials and Methods

### Materials

Design, production and purification of SAP-scFc and AT-03 are described in the supplementary methods and Figure 1. Amyloid-like fibrils of the recombinant λ6 light chain variable domain WIL (rVλ6WIL) were generated by shaking protein (1 mg/mL in PBS) at 37°C and 225 revolutions per min (C24 incubator shaker, New Brunswick Scientific, Edison, NJ) for approximately 72 hours. Fibrils were then purified by centrifugation and stored at 4°C in sterile PBS until used (30). Human amyloid extracts were isolated from amyloid-laden organs using the water floatation method (31).

**Figure 1.**
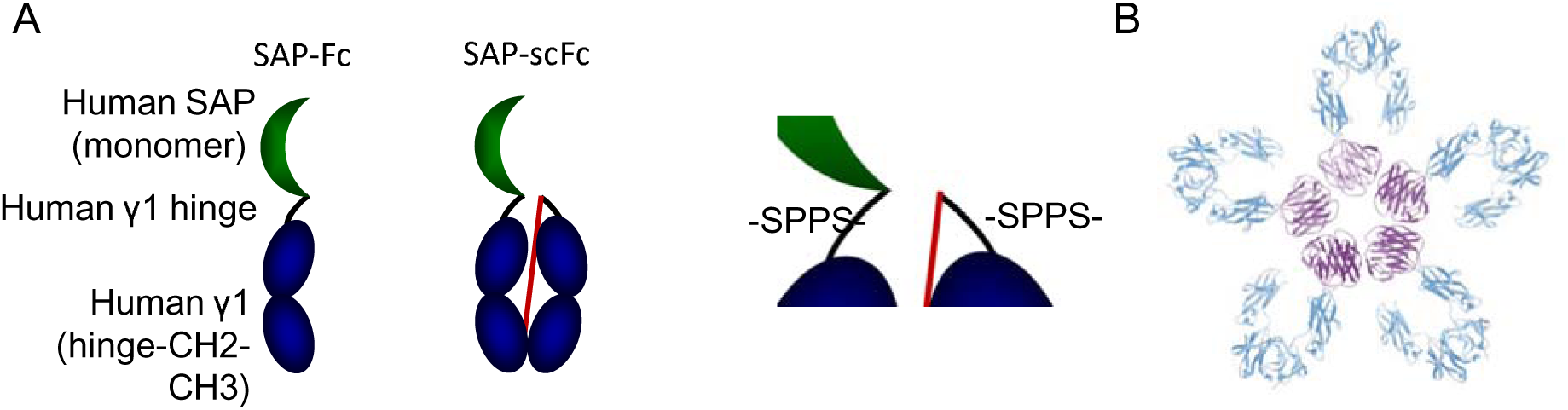
Schematic structure of AT-03 and FcRn binding. (A) A single chain human IgG-Fc fused to SAP comprising one SAP, two CH2, and two CH3 domains. (B) Schematic representation of a AT-03 pentamer where the peripheral blue regions represent human IgG1 Fc and the central purple regions represent SAP.

### Amyloid Clearance in AA mice

AA amyloidosis was induced in immunodeficient VH-LMP2a mice (32) using a single intravenous (IV) injection of amyloid-enhancing factor (AEF) (100 µL) prepared as previously described (31) and a subcutaneous administration of 250 µL of 1% AgNO_3_ on day 0, followed by two further injections on days 7 and 14. Injections of SAP-scFc (3 mg), SAP-Fc (2 mg), human SAP (1 mg), irrelevant (anti-CD20) human IgG1 (6 mg) or PBS were administered at day 21 and the mice euthanized two weeks later. The presence of amyloid in the spleen was assessed histologically using 8-µm thick tissue sections stained with alkaline Congo red (33). Congo red fluorescence was quantified by at least two blinded readers, who scored the amyloid load on a 0–4+ scale. The mean and standard deviation (SD) of the amyloid scores was calculated for each group. Comparisons between groups were performed by Mann-Whitney two-sided test (Prism v8, GraphPad).

### In Vitro Amyloid Binding Studies

Binding of AT-03 to synthetic rVλ6WIL amyloid fibrils and human AL and ATTR amyloid extracts was assessed using a europium-linked immunosorbent assay (EuLISA). The mean and standard deviation of three replicate wells were determined. The data were plotted without the prozone region before fitting to a sigmoid equation with a variable slope (Prism v8, GraphPad) to estimate the EC50. AT-03 protein dilutions were made using the pentamer molecular weight of 410 KDa. AT-03 was biotinylated using Thermo Scientific EZ-Link-Sulfo NHS Biotin (PI21217), according to manufacturer’s instructions, without modification, and stored at 4°C before use. A stock solution was prepared at 2 mg/mL and diluted in Tris-buffered saline with calcium chloride (2 mM) before being used. The binding of biotinylated AT-03 to fresh-frozen human tissues containing AL and ATTR amyloid was assessed by immunohistochemistry following pretreatment with EDTA. Consecutive tissue sections were stained with Congo red (Sigma #C6277). Immunostaining with an anti-human SAP polyclonal antibody (Abcam; #ab45151) was used to demonstrate the distribution of native SAP in tissue sections not treated with EDTA.

### In Vivo Biodistribution Studies

AT-03 protein (50 µg) was radioiodinated with iodine-125 (^125^I) (New England Nuclear, Perkin Elmer) by oxidative coupling using chloramine T (34). The biodistribution of ^125^I-labeled AT-03 was assessed in transgenic mice that constitutively express the human interleukin-6 gene (H2-Ld-IL-6 [H2/IL-6] mice) (35, 36). Systemic AA amyloidosis was induced in these mice by intravenous (IV) injection of 10 µg of AEF. Radioiodinated AT-03 was administered IV (∼100 µCi) and assessed at 48 hours post-injection by small animal SPECT/CT imaging (Siemens Preclinical, Knoxville, TN). Tissue radioactivity (% injected dose per gram) was quantified in 11 organs and tissues harvested at necropsy using a gamma counter. Specific binding of ^125^I-labeled AT-03 with amyloid in the organs was confirmed following micro-autoradiographic and histologic staining of tissue sections with Congo red (37). The AL (38) and AApoAII models are briefly described in the supplementary methods. Twenty four hours after IV injection of 0.5 mg of AT-03, mice were euthanized, and organ samples were snap-frozen and processed for immunofluorescence (IF) studies, as previously described (32). Briefly, the biodistribution of AT-03 in AL and AApoA2 amyloid was evaluated by immunostaining of 8-µm thick tissue sections using an FITC-conjugated anti-human IgG antibody followed by Congo red staining. Mice not injected with AT-03 served as controls.

### Macrophage-Mediated Phagocytosis

Macrophage-mediated phagocytosis of amyloid and amyloid-like fibrils by AT-03 was assessed *in vitro* using monolayers of cultured phorbol myristate acetate (PMA)-activated human THP-1 macrophages. PMA (50 ng/mL; Sigma, P8139) was added and cells allowed to differentiate for 24 hours at 37°C in a 5% carbon dioxide atmosphere. An irrelevant human IgG1-Fc domain served as the negative control. Synthetic rVλ6WIL amyloid-like fibrils or human amyloid extracts were labeled with the pHrodo Red fluorophore (pHrodo™ Red SE, Thermo Fisher, P36600). Fluorescence emission associated with pHrodo red in the macrophages was assessed by fluorescence microscopy (Keyence BZ X800 V 1.3.1). For the single-dose phagocytosis assays, peptides and antibodies were tested at 10 µg. For the dose response phagocytosis assays, peptides and antibodies were tested at 1, 3, 10, and 30 µg. Four fluorescence images were captured for each evaluation. The amount of fluorescence in each image was quantified using image segmentation (Image Pro Premier v. 9.0). The mean and standard deviation of the four images were calculated. Statistical analyses were performed using an unpaired, two-tailed t test with α = 0.05 (Prism v9, GraphPad). Estimates of the Kd (the midpoint concentration of maximal binding) were made by fitting to a single binding site model with specific binding (Prism v9).

### In vivo Macrophage-Mediated Phagocytosis

Human ALλ(SHI) extract was labeled with pHrodo Red STP ester (Life Technologies). Amyloid extract (2 mg), containing 10% (w/w) pHrodo Red-labeled amyloid was injected subcutaneously on the flank of NU/NU mice (*n*=5 per group). Prior to injection, the amyloid extract was incubated with 100 μg of AT-03 or PBS vehicle for 1 h. Serial fluorescence images were acquired under isoflurane anesthesia on days 1, 3, 7, 9, 11 and 14 (iBox Scientia; Analytik Jena, Upland, CA), using a Cy5 excitation and emission filter (100-milisecond exposure, 1 x 1 binning) (39). Endpoint (Day 14) data were analyzed for normality using a Shapiro-Wilk test before comparing using an unpaired Welch’s t-test (α=0.05).

### Ethical Statement

All animal experimental procedures were conducted according to protocols approved by the University of Tennessee, Knoxville Institutional Animal Care and Use Committee (Protocol #1628) or by the French Ministry of Research (APAFIS #7655-2016112211028184 and #33218-2021092417194431). The University of Tennessee animal care and use program holds Association for Assessment and Accreditation of Laboratory Animal Care International (AAALACi) accreditation. The non-human primate study was conducted in accordance with The US Food and Drug Administration (US FDA) Good Laboratory Practice Regulations for Nonclinical Laboratory Studies (GLPs), 21 Code of Federal Regulations (CFR), Part 58 and the applicable principles of the Organisation for Economic Co-operation and Development (OECD) Good Laboratory Practice Principles (GLP) (ENV/MC/CHEM(98)17).

## Results

### Design of AT-03

We took advantage of the known pan-amyloid binding of SAP to create an amyloid-targeting Fc-fusion protein. Two different variants: SAP-Fc were generated comprising from N-term to C-term, the human SAP fused to a single human IgG1-Fc (Hinge-CH2-CH3 regions) and SAP-scFc comprising from N-term to C-term the human SAP fused to an IgG1 Fc (Hinge-CH2-CH3), a (Gly_4_Ser)_3_ linker and a second IgG1 Fc (Hinge-CH2-CH3) (**Figure 1A**). The second variant was designed to allow natural pentamerization of the SAP while keeping an active dimeric Fc domain. To improve the manufacturability and avoid undesirable dimerization of the SAP-scFc protein, codons encoding the cysteine residues in the hinge region of the Fc domain were replaced with those for serine (**Figure 1B**). Fc functionality of the resulting protein, called AT-03, was assessed by binding to human FcRn (**Supplementary Methods**). At pH 6, AT-03 showed comparable binding affinity to human FcRn with Rituximab as a human IgG1 control (**Supplementary Figure 1A**). As expected, both AT-03 and Rituximab demonstrated weak to no binding to human FcRn at pH 7.4 (**Supplementary Figure 1B**).

### Amyloid removal in a murine model of AA amyloidosis

The efficacy of SAP-Fc and SAP-scFc was evaluated in the AgNO_3_ murine model of inducible AA amyloidosis, by monitoring the quantity of amyloid deposits in the spleen. A single IV injection of 3 mg SAP-scFc resulted in a significant reduction in splenic amyloid at 14 days post-injection compared with each of the controls (**Figure 2A and B**). In contrast, SAP-Fc (2 mg) did not significantly reduce amyloid (**Figure 2A**). None of the control agents used at equivalent molar concentration altered the splenic myloid load relative to the phosphate-buffered saline (PBS)–treated control animals. Based on these results, we engineered the AT-03 molecule, derived from the original SAP-scFc scaffold (**Supplementary Methods and Figure 1B**), which was subsequently used for further studies.

**Figure 2.**
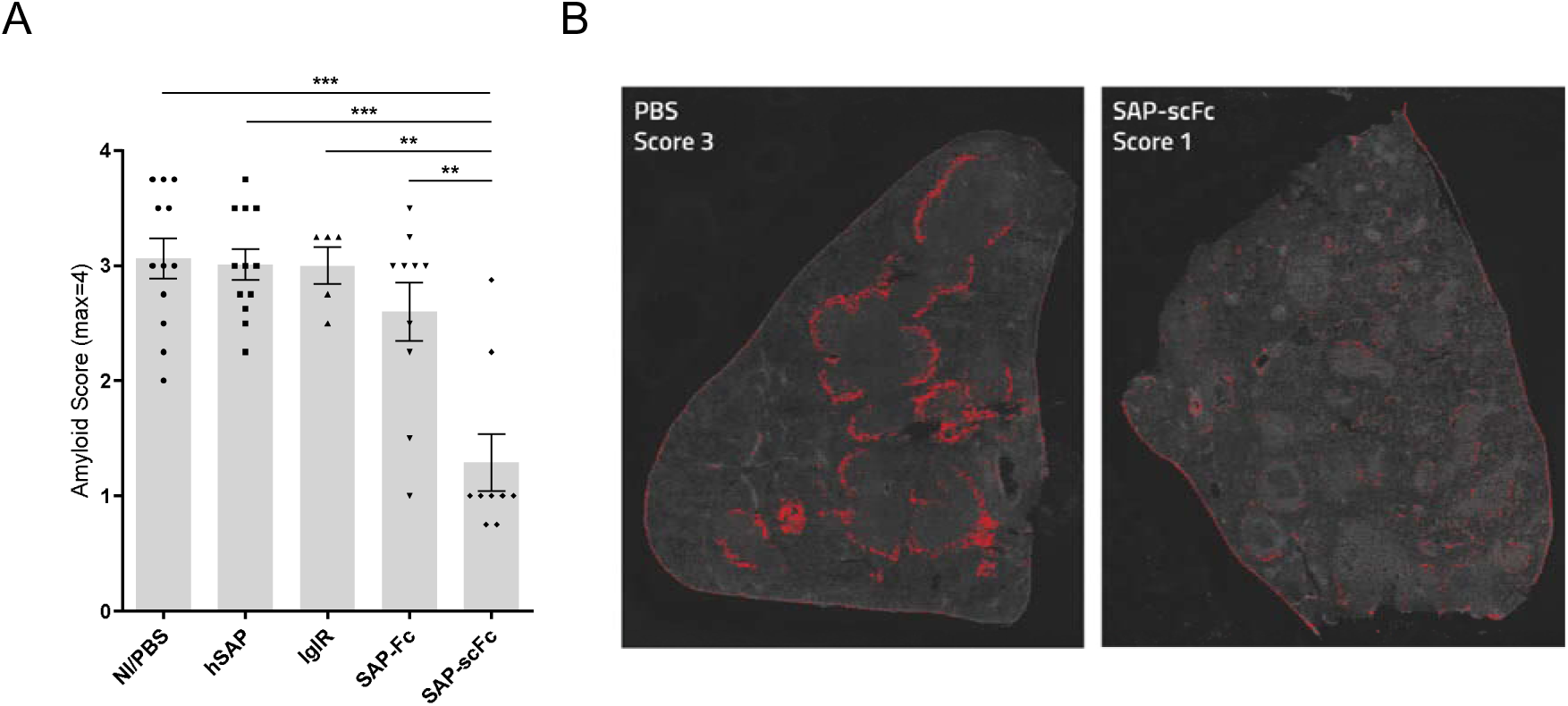
SAP-scFc induces clearance of amyloid deposits in a AA amyloidosis mouse model. (A) SAP-scFc treatment (3 mg) resulted in a significant decrease of splenic AA amyloid load as compared to untreated mice and those treated with human SAP (Hu-SAP, 1mg), non-specific human IgG1 (Rituximab, 6 mg). In contrast, SAP-Fc (2 mg) treatment did not result in amyloid removal. n = 5-12 in 3 independent experiments (except Hu-IgG1, 2 experiments). ** *p* < 0.01, *** p < 0.001, all other comparisons were non significant. (B) Representative fluorescence images of Congo red–stained splenic tissue sections from untreated (PBS) or SAP-scFc-treated mice at 14 days after a single IV administration of reagents.

### AT-03 binds to AL and ATTR amyloid substrates with high potency

The binding of AT-03 was evaluated using synthetic ALλ rVλ6WIL fibrils, as well as human ATTR and AL amyloid extracts by EuLISA. AT-03 demonstrated potent, calcium-dependent, subnanomolar binding, with the estimated half-maximal effective concentration (EC_50_) values ranging from 0.27 nM to 0.63 nM for all amyloid substrates tested (**Figure 3A**). The binding of biotinylated AT-03 to fresh-frozen tissue samples of human AL and ATTR amyloid deposits was assessed by immunohistochemistry, following pre-treatment with EDTA to remove endogenous amyloid-bound SAP (**Figure 3B and C**). The distribution of endogenous SAP in the amyloid-laden tissue sections (without EDTA treatment) was shown to be punctate and discrete and not uniformly distributed throughout the Congo red–birefringent amyloid. Biotinylated AT-03 bound both human AL and ATTR amyloid deposits from the spleen and heart, respectively. Following the EDTA prewash, the binding of biotinylated AT-03 to the tissue was similarly discrete and punctate.

**Figure 3.**
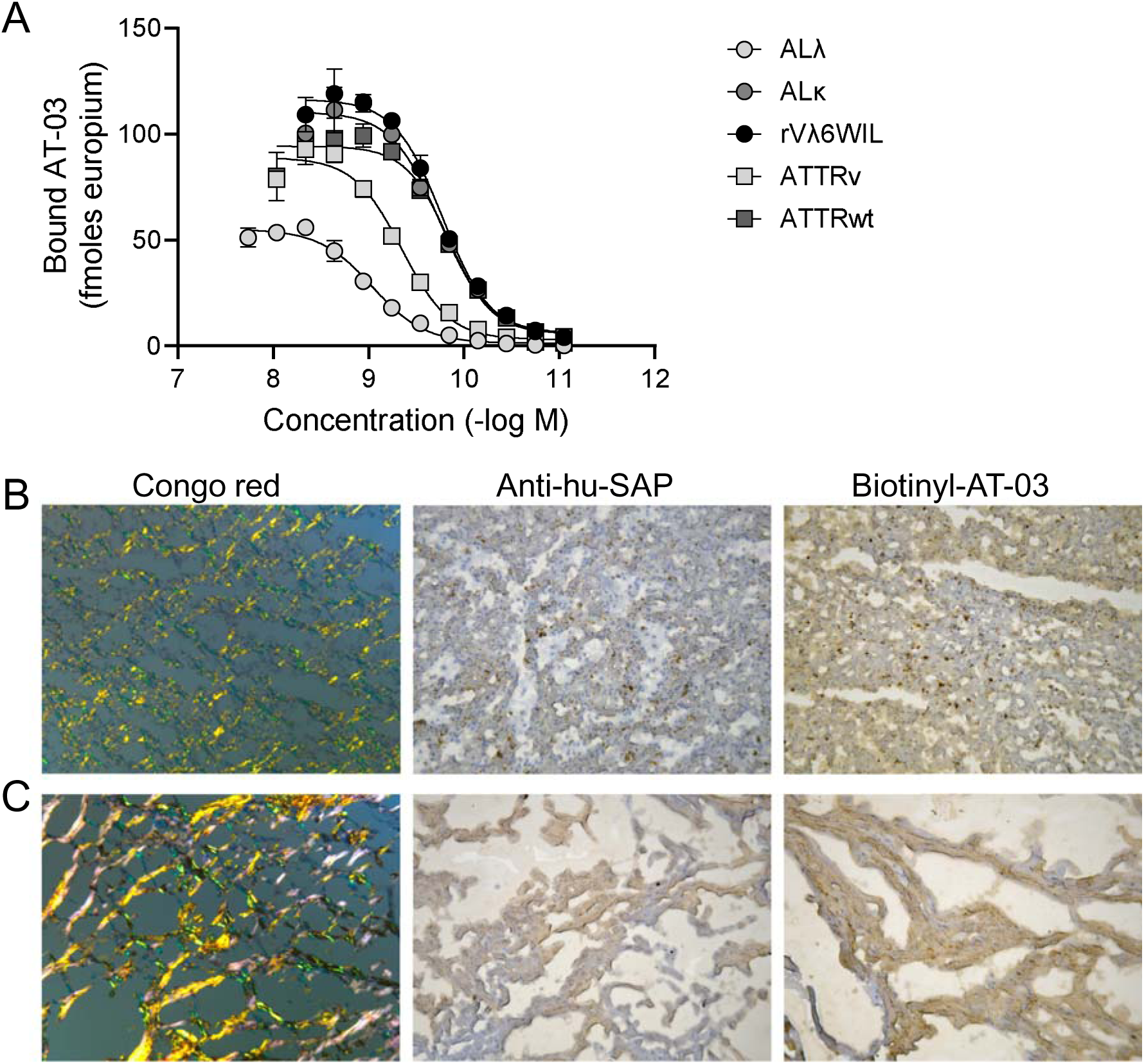
AT-03 binds amyloid with high potency. Binding AT-03 to AL (circle) and ATTR (square) substrates was assessed by EuLISA. Substrates included rVλ6WIL amyloid-like fibrils and ALκ, ALλ, ATTRv, and ATTRwt human amyloid extracts. EC_50_ values were calculated using a 4-parameter logistic regression. Biotinylated AT-03 binds splenic AL (B) and cardiac ATTR (C) amyloid in tissues immunostained using a anti-hu-SAP or biotinylated AT-03 and visualized using diaminobenzidine and compared with Congo red stained sections.

### AT-03 binds specifically to diverse types of amyloid in murine models

The biodistribution of iodine-125-labeled AT-03 (^125^I-AT-03) was assessed in IL-6 transgenic mice with systemic AA amyloidosis using SPECT/CT imaging, tissue biodistribution measurements, and microautoradiography. Intravenous administration of ^125^I-AT-03 (∼10 µg and 124 ± 1.7 µCi) demonstrated visible retention in the liver and spleen of mice with AA amyloidosis at 48 hours post-injection. In contrast, no significant binding was observed in any organ of amyloid-free wild-type (WT) mice (**Figure 4A**). Measurement of tissue-associated radioactivity confirmed significant retention of ^125^I-AT-03 in the liver and spleen of AA mice (5.8% and 9.6% tissue biodistribution (%ID/g), respectively) (**Table 1**). Tissue-to-muscle ratios for the liver and spleen were 36.7 and 60.4, respectively, in AA mice. The AA-to-WT ratio for the liver and spleen was 20 and 17.8 (**Table 1**). In AA mice, the stomach was the only other tissue with >1 %ID/g. Micro-autoradiographic analysis of AA-laden mouse tissues at 48 hours post-injection of ^125^I-AT-03 revealed the presence of radioactivity in the liver, spleen, and heart, which was evident by the presence of black silver grains that were specifically associated with amyloid deposits that also stained positively with Congo red in consecutive tissue sections (**Figure 4B**). Retention of ^125^I-AT-03 in the liver and spleen of AA mice was assessed in a single cohort of mice (*n* = 3) up to 192 hours post-injection following a single IV administration. Serial SPECT/CT imaging of the mice showed visible retention of radioactivity (^125^I-AT-03) at all time points evaluated, up to 192 hours post-injection (**Figure 4C**). Changes in hepatosplenic radioactivity were quantified from the SPECT images (**Figure 4D**). After an initial increase, relative to the 24-hour measurement, splenic radioactivity decreased ∼50% at ∼120 hours post-injection. The hepatic radioactivity decreased linearly from 24 hours post-injection to 192 hours post-injection with a midpoint of ∼100 hours post-injection. The tissue biodistribution of ^125^I-AT-03 was assessed in this cohort of mice at 192 hours post-injection following necropsy (**Supplementary Figure 2A**). At this time, the spleen and liver had retained 2.6 ± 0.4 %ID/g and 1.0 ± 0.5 %ID/g, with the stomach also containing 0.4± 0.4 %ID/g. All other tissues measured were <0.2 %ID/g. ^125^I-AT-03 in the spleen and liver at 192 hours was amyloid-bound based on microautoradiographic evaluation (**Supplementary Figure 2B**). The silver grains seen in the tissue, associated with radioactivity, colocalized with amyloid deposits, which were seen as green-gold deposits in Congo red–stained tissue sections. To better assess the binding of AT-03 to amyloid deposits within cardiac tissue, we used a newly established transgenic mouse model of cardiac AL amyloidosis (38). In these mice, AT-03 (0.5 mg) colocalized within 24 hours post-injection with cardiac amyloid deposits evidenced by immunostaining (**Figure 4E**). Similar colocalization of AT-03 with cardiac, renal, and hepatic amyloid stained with Congo red was observed in a murine model of systemic senile ApoA2–associated amyloidosis (**Figure 4F**).

**Figure 4.**
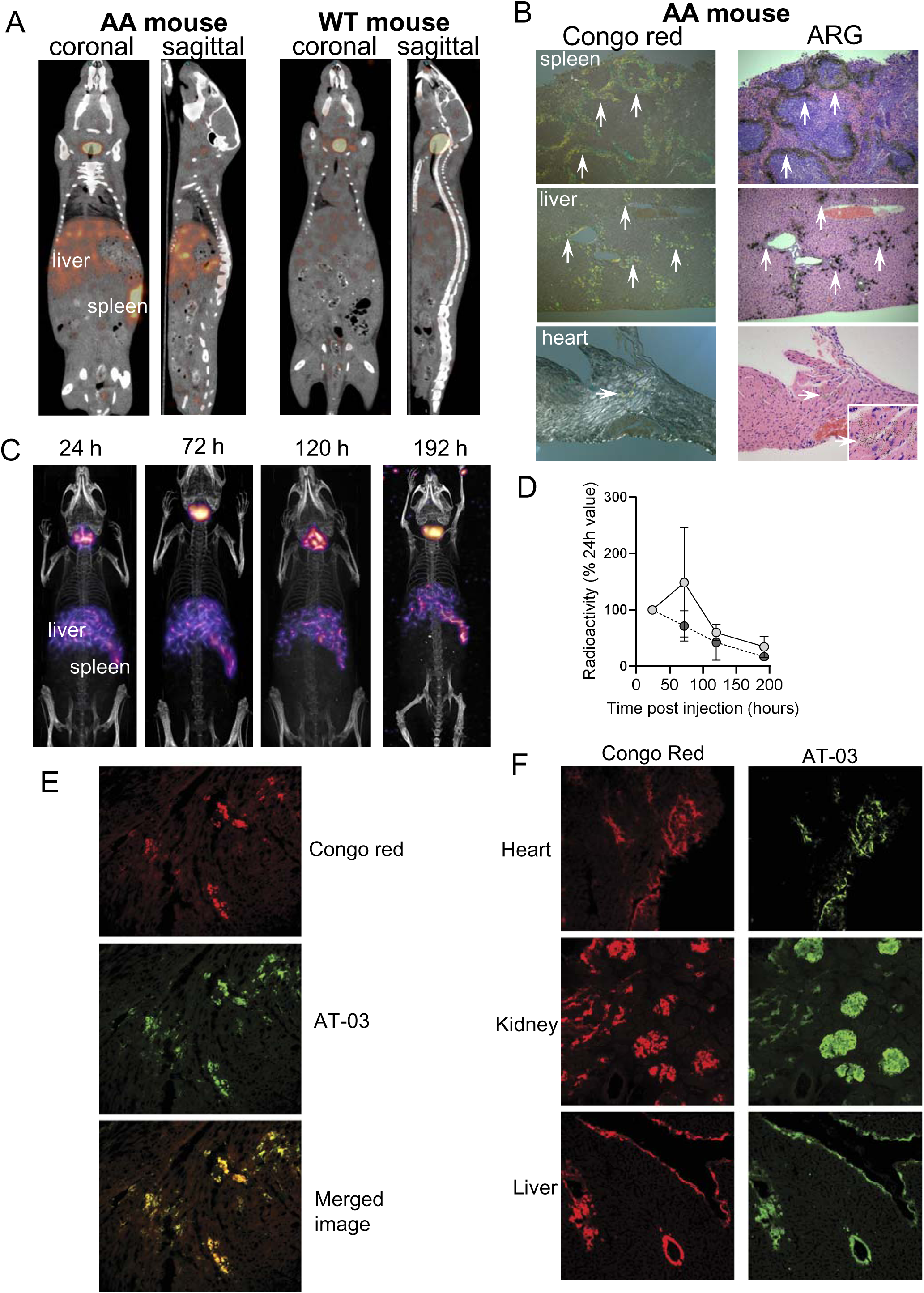
Radioiodinated AT-03 colocalizes specifically with tissue amyloid following IV injection into mice with AA amyloidosis. (A) Small animal SPECT/CT imaging of ^125^I-AT-03 in mice with systemic AA amyloidosis (AA) and amyloid-free wild type (WT) mice at 48 hours post-injection. (B) Specific reactivity with splenic, hepatic, and cardiac amyloid was evidenced using microautoradiography (ARG). (C) Small animal SPECT/CT images (3D coronal view) of AA mice at various time points following IV injection of ^125^I-AT-03. (D) Decrease in splenic (light) and hepatic (dark) radioactivity over 192 hours post-injection, assessed from analysis of SPECT images, following a single IV administration of ^125^I-AT-03 in AA mice. Frozen tissue sections stained obtained from mice were immunostained for the presence of AT-03 in (E) AL (heart) or (F) AApoA2 (heart, kidney, and liver) amyloid deposits using a FITC-conjugated anti-human IgG. The presence of amyloid was shown using Congo red (fluorescence) of consecutive tissue sections.

**Table 1.** Tissue biodistribution (%ID/g) of ^125^I-AT-03 in AA and WT mice at 48 hours post-injection.

| **Organ** | **AA mice** |  | **WT mice** |  | **AA mice**<br>**T:M ratio<sup>4</sup>** | **AA:WT<sup>1</sup>**<br>**ratio** |
| --- | --- | --- | --- | --- | --- | --- |
|  | Mean<br>(%ID/g)<sup>2</sup> | SD<sup>3</sup> | Mean<br>(%ID/g) | SD |  |  |
| Muscle | 0.158 | 0.007 | 0.051 | 0.010 | 1.0 | 3.13 |
| Liver | 5.798 | 2.208 | 0.290 | 0.024 | 36.7 | 19.97 |
| Pancreas | 0.401 | 0.130 | 0.071 | 0.006 | 2.5 | 5.65 |
| Spleen | 9.550 | 4.539 | 0.536 | 0.101 | 60.4 | 17.82 |
| Left kidney | 0.800 | 0.332 | 0.253 | 0.061 | 5.1 | 3.16 |
| Right kidney | 0.629 | 0.331 | 0.278 | 0.058 | 4.0 | 2.26 |
| Stomach | 1.015 | 0.288 | 0.104 | 0.009 | 6.4 | 9.78 |
| Upper intestine | 0.531 | 0.123 | 0.122 | 0.029 | 3.4 | 4.35 |
| Lower intestine | 0.180 | 0.034 | 0.076 | 0.005 | 1.1 | 2.36 |
| Heart | 0.395 | 0.048 | 0.147 | 0.014 | 2.5 | 2.69 |

<sup>1</sup>AA:WT, AA-to-WT mouse organ ratio; <sup>2</sup>%ID/g, percent injected dose per gram of tissue; <sup>3</sup>SD, standard deviation; <sup>4</sup>T:M, tissue-to-muscle ratio.

### AT-03 demonstrated dose-dependent opsonization promoting ex vivo phagocytosis

The impact of AT-03 treatment on the phagocytosis of pHrodo Red–labeled human AL amyloid extract and rVλ6WIL amyloid-like fibrils was assessed *in vitro* using phorbol-myristate acetate (PMA)-activated human THP-1 cells (**Figure 5**). Addition of increasing concentrations of AT-03 (6 nM, 20 nM, 60 nM, and 200 nM) to rVλ6WIL, human ALκ, or AL λ amyloid extracts enhanced phagocytosis in a dose dependent manner, which reached a maximum effect at ∼ 60 nM (**Figure 5A**). Addition of 20% v/v human serum (HS) as a source of complement factors significantly enhanced the uptake of the substrates in the presence of 60 nM AT-03 (**Figure 5B**). AT-03–mediated enhancement of phagocytosis of human ALλ amyloid extract was further assessed using an *in vivo* amyloidoma model and monitored using optical imaging (**Figure 5C and 5D**). pHrodo Red–labeled ALλ amyloid extract (2 mg) was pre-incubated with AT-03 (100 µg) or PBS before being implanted subcutaneously in NU/NU mice (*n*=5 per group). The pHrodo red fluorescence emission from the amyloid mass at 14 days post-implantation was significantly enhanced in mice receiving AT-03–treated amyloid extract as compared to untreated control mice (*p*=0.0098) (**Figure 5C and D**).

**Figure 5.**
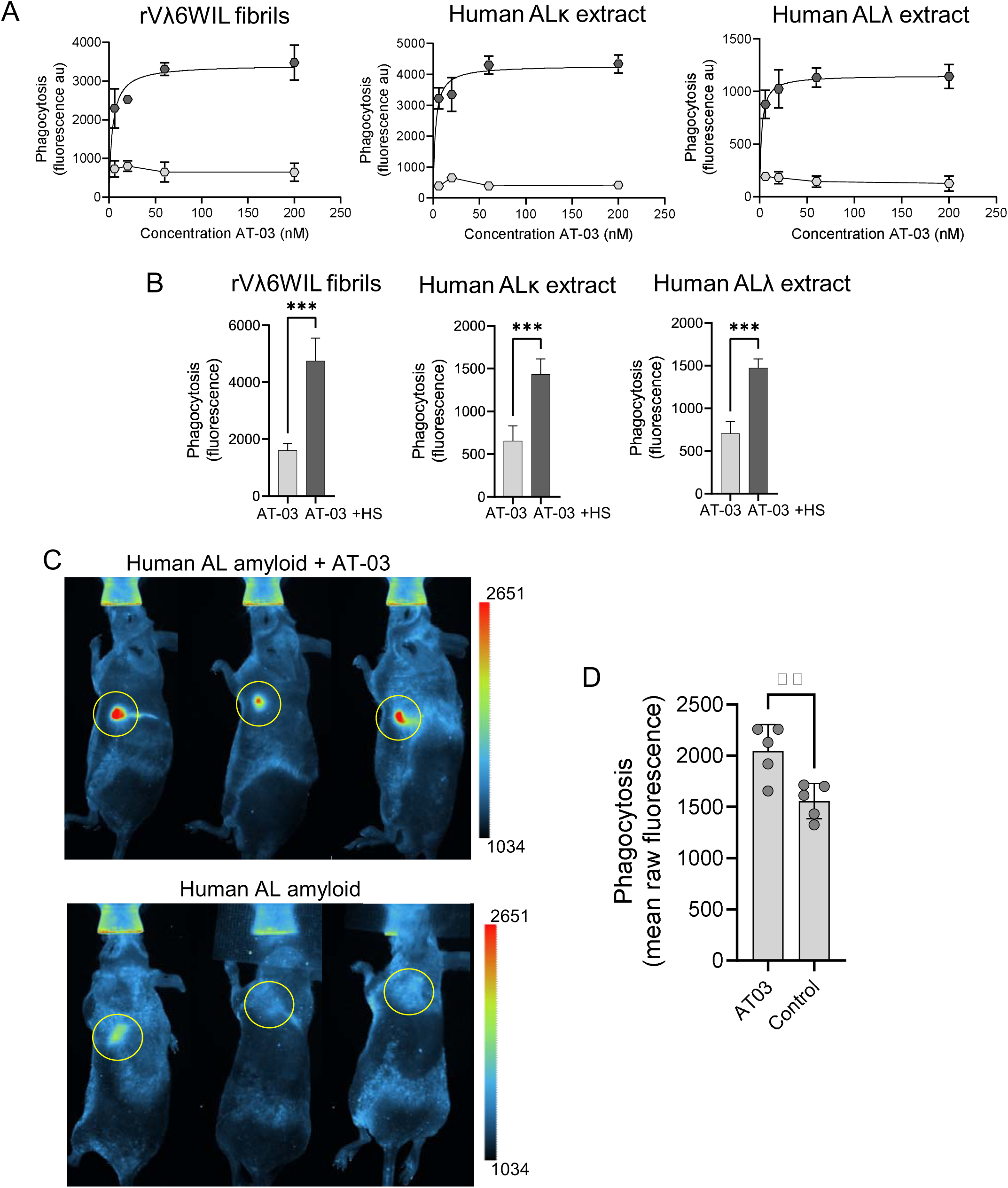
Phagocytosis of human AL amyloid extracts is enhanced by opsonization by AT-03 and human complement. (A) Phagocytosis of pHrodo Red–labeled synthetic rVλ6WIL fibrils, or human ALκ, and ALλ amyloid extracts was enhanced with increasing doses of AT-03 (dark) as compared to the vehicle control (light). (B) The addition of human serum (20% v/v; HS) significantly increased AT-03–induced (60 nM) phagocytosis of the substrates. Different batches of pHrodo red-labeled subtrates were used for studies in A and B, which explains the different intensity in the AT-03 fluorescence at 60 nM. (C) Pre-treatment of pHrodo Red–labeled human ALλ amyloid extract with AT-03 (top) increased *in vivo* phagocytosis, which was evident by difference in pHrodo Red fluorescence emission of the amyloid mass (circle) detected by optical imaging (representative images acquired at day 14 post injection; *n*=5 per group) compared to human ALλ amyloid extracts alone (bottom). (D) Change in phagocytosis of amyloid-bound pHrodo Red at Day 14 following subcutaneous injection in mice. Human AL amyloid extract was either treated with AT-03 (dark) or untreated (light).

### AT-03 pharmacokinetics in a non-human primate model

In cynomolgus monkeys, IV administration of AT-03 at 3 mg/Kg or 10 mg/Kg resulted in mean maximum serum concentration (Cmax) values of 69.8 µg/mL and 199 µg/mL, respectively. Exposure, measured using area under the curve analysis (AUCinf), increased proportionally from 732 h*µg/mL to 2060 h*µg/mL in the 3 mg/Kg and 10 mg/Kg-treated animals. No notable differences observed between sexes. All serum dilution concentrations of AT-03 remained above the assay lower limit of quantitation (LLOQ) of 5 ng/mL, through the final 336-hour time point. At a dose of 3 mg/Kg, the mean elimination half-life (*t*_1/2_) of AT-03 was 76.3 hours increasing marginally to 83.8 h for the 10 mg/Kg dose (**Supplementary Figure 3**).

## Discussion

Despite recent advances in reducing or stabilizing amyloid proteins, systemic amyloidosis remains frequently fatal, as substantial organ deposition is often present at diagnosis (20, 21). There is therefore an urgent need for therapies that actively remove amyloid and rapidly restore organ function. Taking advantage of the specific binding of SAP to all types of amyloid, AT-03, a pan-amyloid binding fusion protein comprising the amyloid reactivity of SAP and the Fcγ1 domain for immune cell activation, has been developed. *In vitro*, AT-03 binds potently to AL and ATTR human amyloid extracts and *in vivo*, to amyloid deposits in the liver, kidney, spleen, but also in heart in multiple mouse models, including AL-, AA-, and ApoA2-associated amyloidosis. Furthermore, significant removal of splenic amyloid deposits was demonstrated in a mouse model of AA amyloidosis following a single 3-mg IV dose of SAP-scFc, a prototypic structurally homologous precursor of AT-03.

AT-03 demonstrated dose-dependent induction of *in vitro* phagocytosis of human AL amyloid extracts by PMA-activated human THP-1 cells, which was further enhanced by the presence of human serum as a source of complement. The importance of complement-mediated phagocytosis in amyloid removal has previously been demonstrated *in vivo* with anti-SAP antibodies (40, 41). The therapeutic potential of AT-03 was further evaluated in a murine AL amyloidoma model, in which pre-treatment of the amyloid with 100 µg of AT-03 enhanced *in vivo* phagocytosis. Unfortunately, because the new AL amyloidosis transgenic model lacks full penetrance and reliable biomarkers of amyloid burden, it could not be used to evaluate AT-03 in a more physiological setting. SPECT/CT imaging demonstrated uptake of ^125^I-AT-03 in AA amyloid laden abdominothoracic organs of mice, including liver and spleen, but not in the heart (25–27). We speculate that the scant deposits of cardiac AA amyloid, and the inherent resolution of the imaging platform may account for the inability to observe the uptake of ^125^I-AT-03 in heart but we cannot formally exclude a limited access to the myocardium as previously shown with SAP imaging (29). However, AT-03 was readily detected in association with cardiac amyloid deposits both in AA, as shown by microautoradiographs following ^125^I-AT-03 injection and in transgenic AL and APOA2 mice after IV administration of AT-03, demonstrating its ability to penetrate the myocardium of mice and bind amyloid therein. Murine amyloidosis models lack human SAP as an endogenous competitor for AT-03 binding; thus, therapeutic efficacy for cardiac amyloid removal should be further assessed in pre-clinical models expressing human SAP.

Currently, no approved therapy is designed to remove amyloid deposits in systemic amyloidosis. Several investigational monoclonal antibodies have shown potential to clear amyloid and improve organ function, but results remain mixed: Dezamizumab, a fully humanized anti-SAP mAb, showed the potential for immune-mediated amyloid clearance (12, 28) but was discontinued after a serious adverse event (29). Birtamimab, a humanized mAb that binds a cryptic epitope exposed on misfolded light chain showed early benefit (42) but failed in phase 3. Anselamimab induced organ responses in phase 1 (43) but did not meet endpoints in phase 3 except in a specific subset. ALXN2220, an anti-misfolded TTR reduced cardiac amyloid in ATTR in a phase 1b trial (44) and is now in phase 3. None of these antibodies, however, display pan-amyloid reactivity which supports our approach toward developing a fusion protein with the potential for pan-amyloid removal.

In conclusion, there remains an unmet need for therapies that safely remove amyloid deposits and restore organ function in systemic amyloidosis. AT-03 showed highly specific binding to human and synthetic amyloid fibrils, enabled targeting across multiple organs in murine models, enhanced macrophage-mediated phagocytosis *ex vivo* and *in vivo* in an amyloidoma model and reduced splenic deposits in AA amyloidosis. While in vivo efficacy in other models, particularly with cardiac involvement, remains to be confirmed, these findings support the evaluation of AT-03 as a potential therapeutic for systemic amyloidosis.

## Acknowledgments

We appreciate the assistance of Jim Wesley in the preparation of the microautoradiographs and Daniel C. Wooliver for histological studies. We thank Aaron Endsley (Certara) for his work analyzing PK data from the non-human primate (NHP) studies and the staff of the Biologie Intégrative Santé Chimie Environnement (BISCEm) technical platforms at the University of Limoges. Medical writing assistance for an early draft was provided by Bridget Healy, MBChB, MPH, and David Gibson, PhD, CMPP, of ApotheCom and was funded by Attralus, Inc.

## Author Contributions

Conceptualization, C.S., A.J., F.B., S.G., S.J.K., and J.S.W.; methodology, C.S., G.R.C., M.C., F.B., G.B., S.G., S.S., S.J.K., A.D.W., J.S.F., and J.S.W.; validation, S.S.; investigation, C.S., G.R.C., A.J., S.B., G.M.R., F.B., M.C., , M.C., C.V., C.C., G.B., S.G., S.S., S.J.K, A.D.W., T.R., A.S., M.B., J.S.F., and J.S.W.; writing—original draft preparation, C.S., A.J., S.B., F.B., G.B., S.G., S.S., and J.S.W; writing—review and editing, C.S., A.J., S.B., F.B., S.J.K., E.B.M., J.S.F., and J.S.W.; visualization, C.S. and J.S.W; supervision, C.S., and J.S.W.; project administration, C.S., and J.S.W.; funding acquisition, C.S., S.S., G.B., and J.S.W. All authors have read and agreed to the published version of the manuscript.

## Abbreviations

mAb: monoclonal antibody
Fc: Fragment crystallizable
SAP: serum amyloid P component
IV: intravenous
EuLISA: europium-linked immunosorbent assay
IF: immunofluorescence
PMA: phorbol myristate acetate

