## Supplementary information for "Preclinical characterization of AT-03, a novel Serum Amyloid P fusion protein that demonstrates pan-amyloid binding and removal"

- **Supplementary methods**
- **Supplementary references**
- **Supplementary figures**
  - **Supp Figure S1**
  - **Supp Figure S2**
  - **Supp Figure S3**

**Supplementary methods:**

**Production and purification of proteins**

SAP-scFc, SAP-Fc and AT-03 were produced in CHO, purified by protein A affinity and anion exchange chromatography and treated to remove endotoxins. AT-03 was further processed through a solvent/detergent viral inactivation step and viral filtration step. Purified human SAP was kindly provided by the Laboratoire Français du Fractionnement et des Biotechnologies (LFB) and the human IgG1 used as control was obtained from Rituximab stock leftovers.

**Mouse models of AL and AApoAII amyloidosis.**

The mouse model for AL amyloidosis is described in (32). Briefly, it uses a double knock-in strategy based on the targeted insertion into the Ig kappa locus of a gene coding a human Ig light chain from a patient with AL amyloidosis (λS). This allows the overexpression, by plasma cells, of a human AL light chain. To produce only free LC, we took advantage of the inactivation of the IgH locus by the latent membrane protein 2A gene from the Epstein-Barr virus allowing for a normal plasma cell development without Ig heavy chain expression. AL amyloidosis in this model was induced by a single 200-µg injection of preformed AL amyloid-like fibrils, generated *in vitro* from purified λS VL domains. AL amyloid deposition appeared at 2 to 6 months post-injection in approximately two third of mice, with predominantly cardiac amyloid deposits. For other organs, we used plasminogen activator inhibitor (PAI-1) knockout mice under a C57/Bl6 genetic background that exhibit accelerated senile ApoA2 amyloidosis [1]. These mice develop amyloidosis starting at 12 months of age, with deposits in kidney, spleen and, to a lesser extent, liver and heart.

**Binding to Human FcRn**

The binding of AT-03 to human FcRn was performed using a Biacore 8K (Cytiva), with a basic set-up of a CM5 chip (Cytiva) and a running buffer of PBS with Tween at pH 6.0 or pH 7.4. AT-03 was immobilized with a contact time of 30 seconds, flow rate of 10 µL/min, and a running buffer of HBS-EP+ (10 mM HEPES, 0.15 M NaCl, 3 mM EDTA and 0.05% v/v Surfactant P20). Working concentrations of human FcRn of 46.9, 93.8, 187.5, 375, 750, 1500, 3000, and 6000 nM had an association time of 60 seconds, dissociation time of 90 seconds, and flow rate of 30 µL/min, followed by addition of regeneration buffer of PBS (pH 7.4). The binding affinity was evaluated by fitting to a steady-state affinity model using Biacore evaluation software (Cytiva).

**Pharmacokinetic Studies in a Non-Human Primate**

The pharmacokinetic analysis of AT-03 was evaluated following a single 3 mg/Kg or 10 mg/Kg IV administration of AT-03 in male and female cynomolgus monkeys. Serum samples were collected at prespecified time points up to 336 hours post-injection. The concentration of AT-03 at each time point was estimated using an EuLISA system, with the mean and standard deviation for each concentration of standard. Pharmacokinetic analyses were performed on AT-03 concentration data using noncompartmental analysis (Phoenix WinNonlin, Version 8.3).
