## Supplementary Figures 1-3 for "Preclinical characterization of AT-03, a novel Serum Amyloid P fusion protein that demonstrates pan-amyloid binding and removal"

### Slide 1
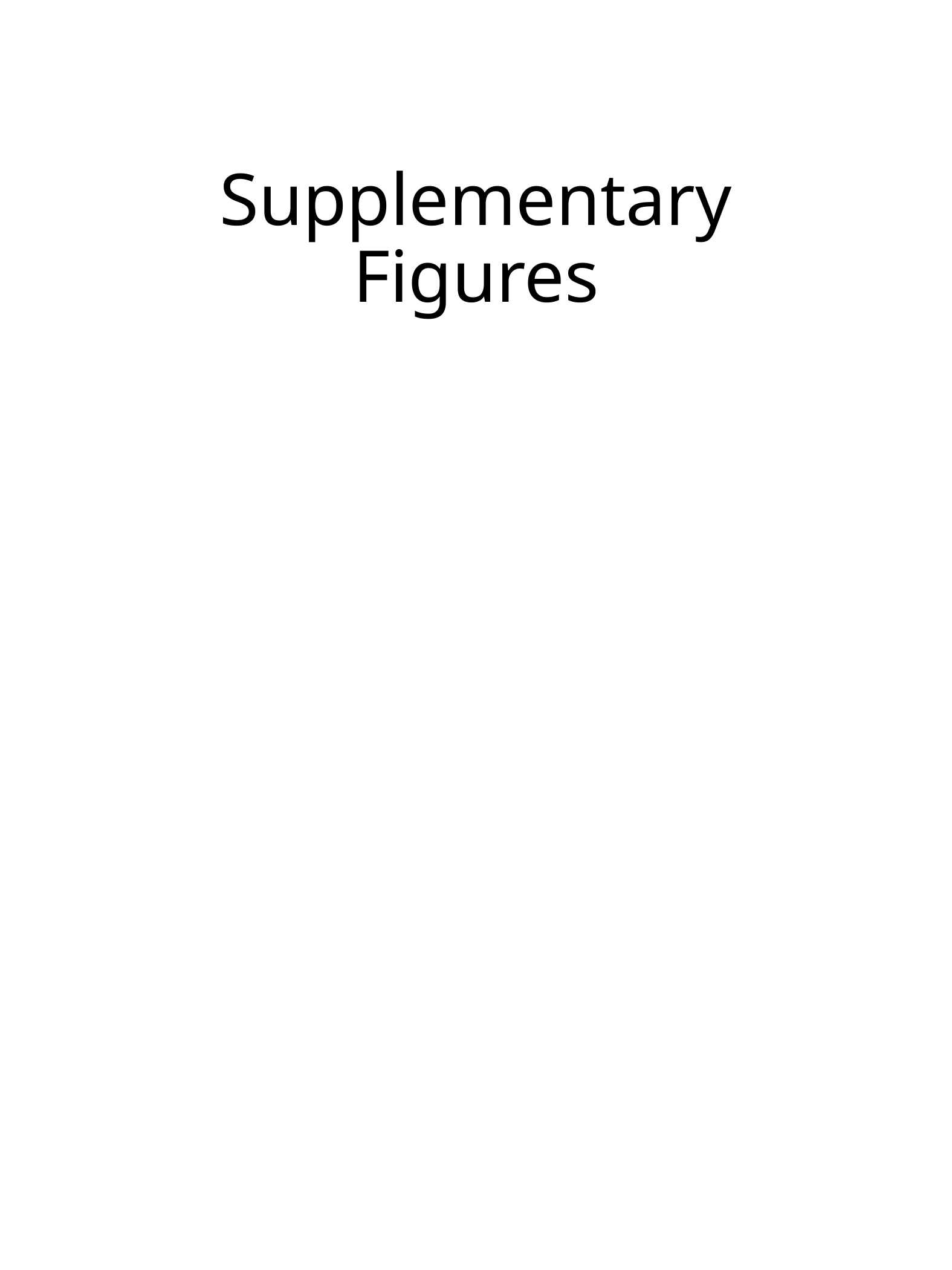

Supplementary Figures

### Slide 2
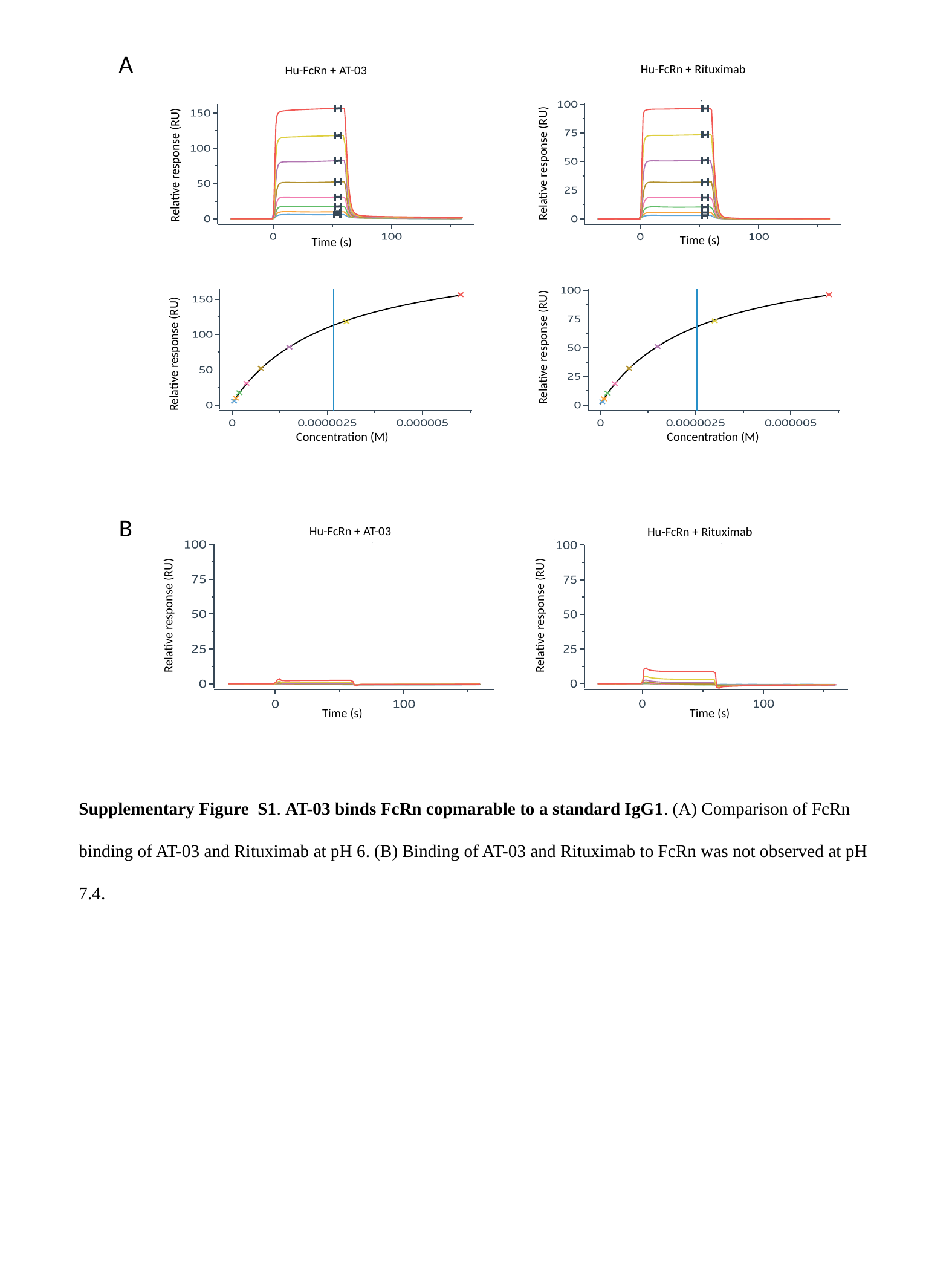

A
Hu-FcRn + Rituximab
Hu-FcRn + AT-03
Relative response (RU)
Relative response (RU)
Time (s)
Time (s)
Relative response (RU)
Relative response (RU)
Concentration (M)
Concentration (M)
B
Hu-FcRn + AT-03
Hu-FcRn + Rituximab
Relative response (RU)
Relative response (RU)
Time (s)
Time (s)
Supplementary Figure S1. AT-03 binds FcRn copmarable to a standard IgG1. (A) Comparison of FcRn binding of AT-03 and Rituximab at pH 6. (B) Binding of AT-03 and Rituximab to FcRn was not observed at pH 7.4.

### Slide 3
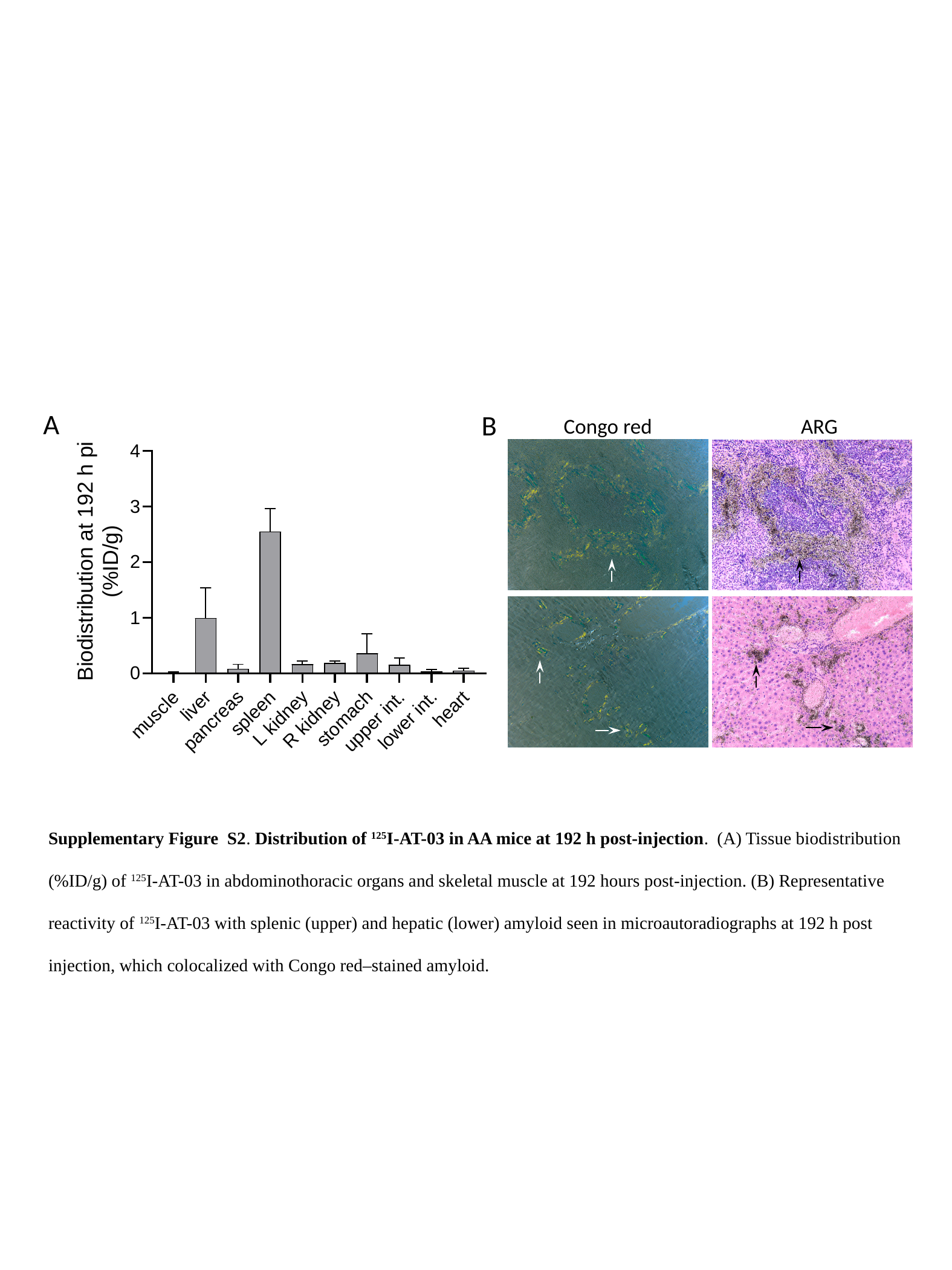

A
B
Congo red
ARG
Supplementary Figure S2. Distribution of ^125^I-AT-03 in AA mice at 192 h post-injection. (A) Tissue biodistribution (%ID/g) of ^125^I-AT-03 in abdominothoracic organs and skeletal muscle at 192 hours post-injection. (B) Representative reactivity of ^125^I-AT-03 with splenic (upper) and hepatic (lower) amyloid seen in microautoradiographs at 192 h post injection, which colocalized with Congo red–stained amyloid.

### Slide 4
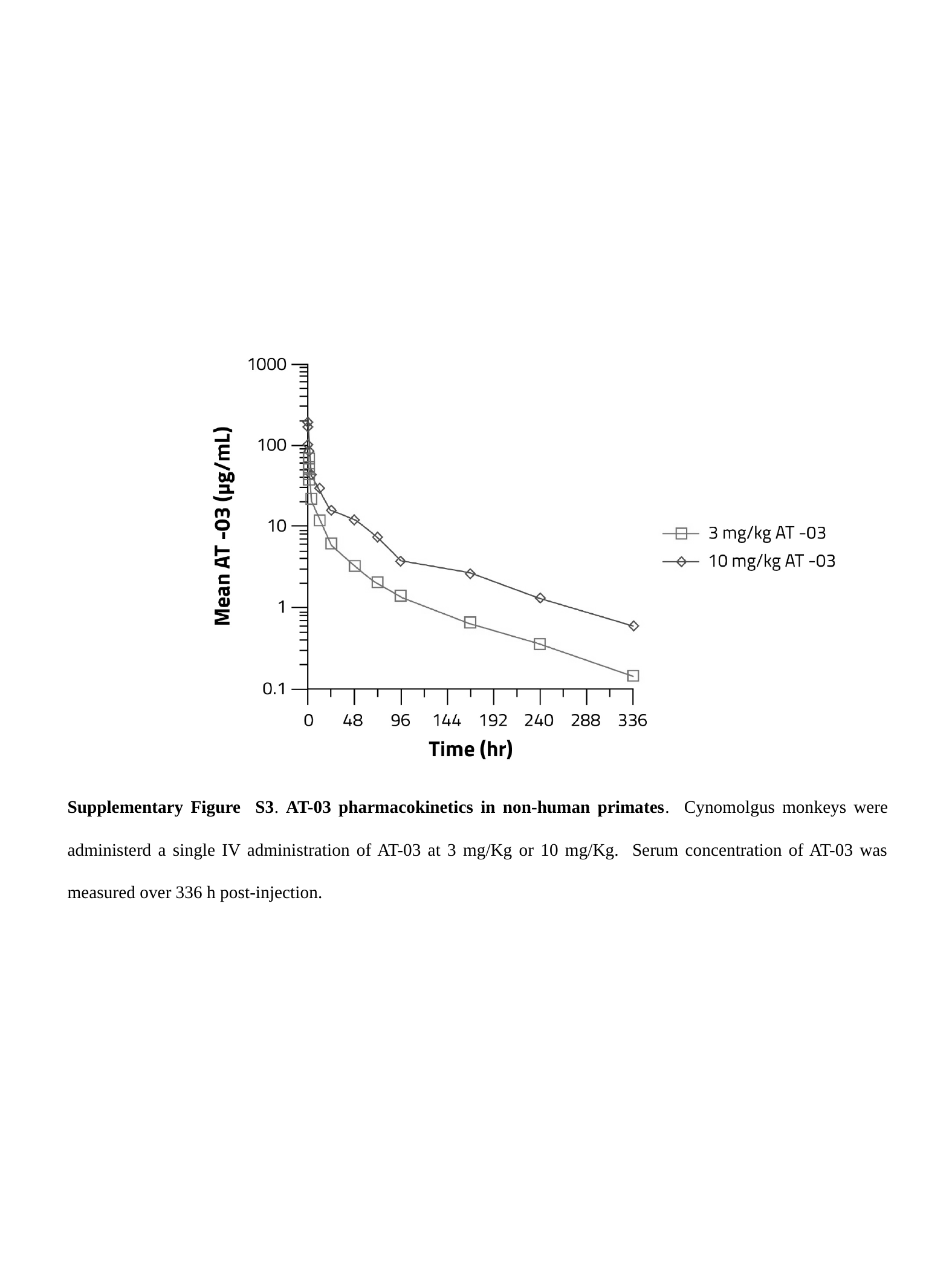

Supplementary Figure S3. AT-03 pharmacokinetics in non-human primates. Cynomolgus monkeys were administerd a single IV administration of AT-03 at 3 mg/Kg or 10 mg/Kg. Serum concentration of AT-03 was measured over 336 h post-injection.
